# Meta-analysis of genome-wide association studies for body fat distribution in 694,649 individuals of European ancestry

**DOI:** 10.1101/304030

**Authors:** Sara L. Pulit, Charli Stoneman, Andrew P. Morris, Andrew R. Wood, Craig A. Glastonbury, Jessica Tyrrell, Loïc Yengo, Teresa Ferreira, Eirini Marouli, Yingjie Ji, Jian Yang, Samuel Jones, Robin Beaumont, Damien C. Croteau-Chonka, Thomas W. Winkler, Giant Consortium, Andrew. T. Hattersley, Ruth J. F. Loos, Joel N. Hirschhorn, Peter M. Visscher, Timothy M. Frayling, Craig A. Glastonbury, Hanieh Yaghootkar, Cecilia M. Lindgren

## Abstract

One in four adults worldwide are either overweight or obese. Epidemiological studies indicate that the location and distribution of excess fat, rather than general adiposity, is most informative for predicting risk of obesity sequellae, including cardiometabolic disease and cancer. We performed a genome-wide association study meta-analysis of body fat distribution, measured by waist-to-hip ratio adjusted for BMI (WHRadjBMI), and identified 463 signals in 346 loci. Heritability and variant effects were generally stronger in women than men, and we found approximately one-third of all signals to be sexually dimorphic. The 5% of individuals carrying the most WHRadjBMI-increasing alleles were 1.62 times more likely than the bottom 5% to have a WHR above the thresholds used for metabolic syndrome. These data, made publicly available, will inform the biology of body fat distribution and its relationship with disease.

## Introduction

One in four adults worldwide are either overweight or obese (1, 2) and are at increased risk of metabolic disease. While higher adiposity increases morbidity and mortality (1, 3), epidemiological studies indicate that the location and distribution of excess fat within particular depots is more informative than general adiposity for predicting disease risk. Independent of their overall body mass index (BMI), individuals with higher central adiposity have increased risk of cardiometabolic diseases, including type 2 diabetes (T2D) and stroke (4, 5); in contrast, individuals with higher gluteal adiposity have lower risk of such outcomes.(5) Previous studies indicate that fat distribution, as assessed by waist-to-hip ratio (WHR), is a trait with a strong heritable component, independent of overall adiposity (measured by BMI), with twin-based heritability estimates ranging between 30-60% (5, 6) and narrow-sense heritability estimates have been estimated at ~50% in women and ~20% in men (5). The most recent genome-wide association study in 224,459 samples implicated 49 loci associated with WHR adjusted for BMI (5), and recent Mendelian randomisation studies using known WHR-associated genetic variants showed putative causal effects of higher WHR on T2D and coronary artery disease independently of BMI (7).

## Results

With the goal of pinpointing genetic variants associated to body shape and fat distribution and motivated by the recent release of genetic data from half a million individuals (8), we performed a meta-analysis of WHR adjusted for BMI (WHRadjBMI). WHRadjBMI is an easily-measured fat distribution phenotype that correlates well with imaging-based fat distribution measures (9). We performed genome-wide association studies (GWAS) of WHRadjBMI in the UK Biobank data set (8), a collection of 484,563 samples with densely-imputed genotype data, using a linear mixed model (10) to account for relatedness and ancestral heterogeneity. We then combined the results with publicly-available GWAS data generated by the GIANT consortium for the same phenotype (**Table 1** and **Methods**) (5), resulting in a meta-analysis of 694,649 samples (**Table 1**) and ~27.4M SNPs (**Methods**). As a sensitivity analysis and to evaluate the robustness of our results, we also performed a GWAS of WHR unadjusted for BMI (**Table 1**).

**Table 1.**
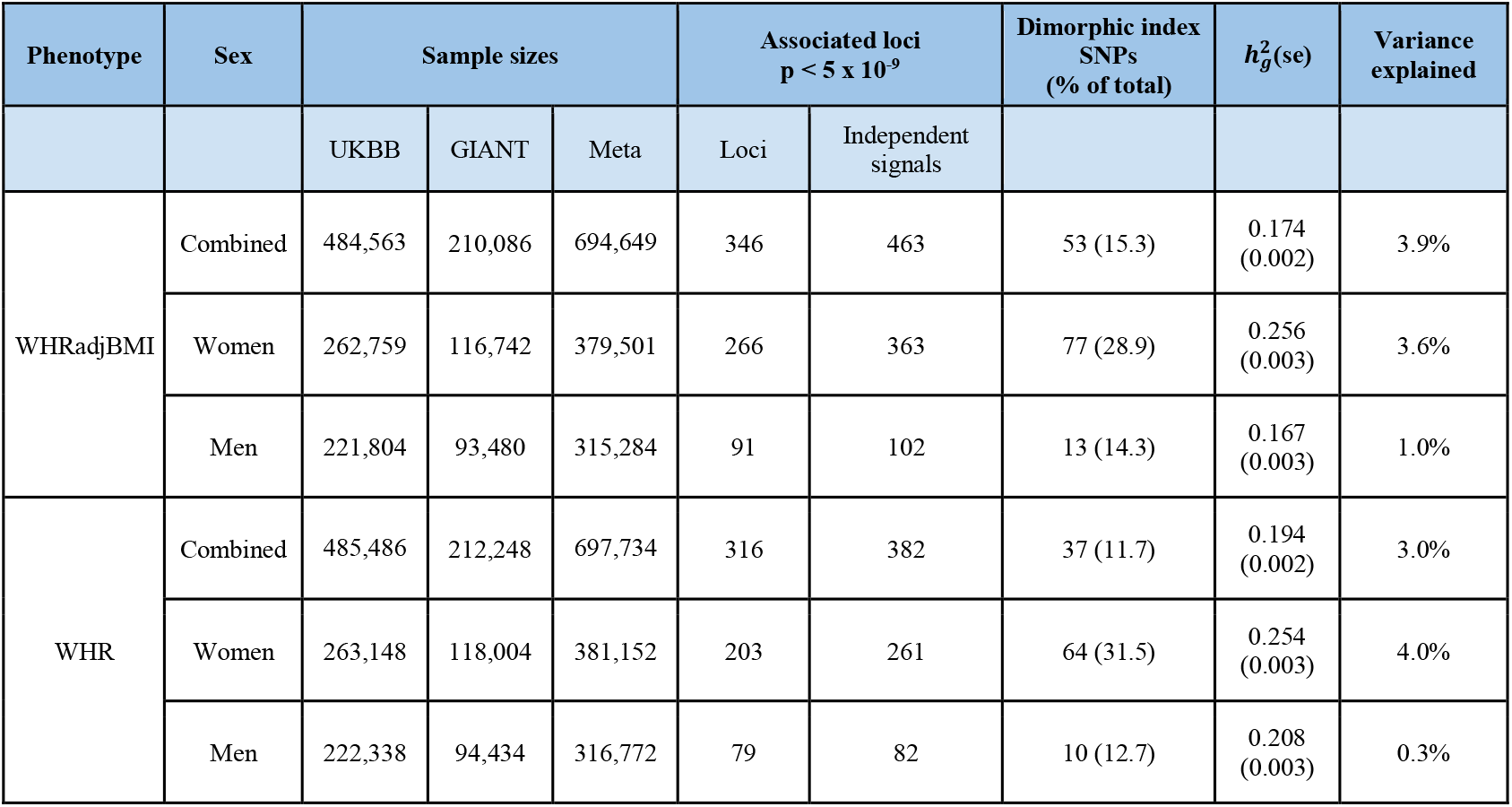
| Large-scale meta-analysis in body fat distribution. We performed a meta-analysis of fat distribution as measured by WHRadjBMI in up to 694,649 individuals. We performed analyses of WHR as a sensitivity measure. Our analyses increase the number of WHRadjBMI-associated loci (p < 5 × 10^−9^, to account for SNP density in UK Biobank) to 346 loci. SNP-based heritability 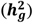 results, estimated using the restricted maximum likelihood method implemented (10), and top-associated loci indicate patterns of sex-dimorphism. The top-associated index SNPs explain 3.9% of the overall phenotypic variance (i.e., adjusted R^2^) in fat distribution (calculated in an independent dataset, N = 7,721).

| Phenotype | Sex | Sample sizes | | | Associated loci<br>$p < 5 \times 10^{-9}$ | | Dimorphic index<br>SNPs<br>(% of total) | $h_g^2$(se) | Variance explained |
| --- | --- | --- | --- | --- | --- | --- | --- | --- | --- |
|  |  | UKBB | GIANT | Meta | Loci | Independent signals |  |  |  |
| WHRadjBMI | Combined | 484,563 | 210,086 | 694,649 | 346 | 463 | 53 (15.3) | 0.174 (0.002) | 3.9% |
|  | Women | 262,759 | 116,742 | 379,501 | 266 | 363 | 77 (28.9) | 0.256 (0.003) | 3.6% |
|  | Men | 221,804 | 93,480 | 315,284 | 91 | 102 | 13 (14.3) | 0.167 (0.003) | 1.0% |
| WHR | Combined | 485,486 | 212,248 | 697,734 | 316 | 382 | 37 (11.7) | 0.194 (0.002) | 3.0% |
|  | Women | 263,148 | 118,004 | 381,152 | 203 | 261 | 64 (31.5) | 0.254 (0.003) | 4.0% |
|  | Men | 222,338 | 94,434 | 316,772 | 79 | 82 | 10 (12.7) | 0.208 (0.003) | 0.3% |

We identified 346 loci (300 novel) containing 463 independent signals associated with WHRadjBMI (p < 5 × 10^−9^, to account for the denser imputation data (11); **Methods, Supplementary Table 1** and **Supplementary Fig 1**). The Linkage Disequilibrium (LD) Score Regression (12) intercept (1.035) of the meta-analysis results indicated that the observed enrichment in genomic signal was due to polygenicity and not confounding (**Supplementary Table 2**). Of the 300 novel signals, 234 (78%, p_binoimal_ < 1 × 10^−7^) were directionally-consistent in an independent dataset with a relatively small sample size (N = 7,721) and signals were consistent in several sensitivity checks (**Supplementary Tables 3-5**, and **Supplementary Fig 2-3**). Combined, these variants explained ~3.9% of the variance in WHRadjBMI in the independent study (**Methods** and **Table 1**). We constructed a weighted polygenic risk score using the 346 index SNPs discovered in the combined meta-analysis and tested this score in the same independent study. The 5% of individuals carrying the most WHRadjBMI-raising alleles were 1.62 times more likely to meet the WHR threshold used to define metabolic syndrome (13) than the 5% carrying the fewest (consistent with the results obtained from unweighted polygenic score; **Methods**). The WHRadjBMI of people in the top 5% of the PRS was 1.05 and 1.06 times greater in men and women, respectively, compared to those in the bottom 5% of the PRS.

To investigate the potential for collider bias resulting from conditioning WHR on BMI, we investigated the behavior of WHRadjBMI-associated SNPs in GWAS of WHR (without adjustment for BMI) and BMI alone. We found that the majority of WHRadjBMI signals identified have genuine effect on body shape, and that any bias caused by adjusting WHR for a correlated covariate (14, 15) (that is, BMI) was minimal. Of the 346 index variants, 311 associated with stronger standard deviation effect sizes for WHR (unadjusted) than with standard deviation effect sizes for BMI (**Supplementary Table 3** and **Supplementary Fig 4**). This observation also indicates that the WHR association is unlikely to be secondary to the known effect of higher BMI resulting in higher WHR. Furthermore, the common SNP associated with the largest known effect on BMI, that in the *FTO* gene (16), was not associated with WHRadjBMI (rs1421085, p = 0.40) despite a very strong association with WHR (p = 4 × 10^−118)^). Finally, carrying each additional (weighted) WHRadjBMI-raising allele was associated with an increase in WHRadjBMI of 0.0199 SD (p = 6 × 10^−62^; adjusted R^2^ = 4%), an increase in WHR of 0.0111 SD (p = 3 × 10^−20^; adjusted R^2^ = 0.12%) and a decrease in BMI of 0.0038 SD (p = 1.4 × 10^−3^; adjusted R^2^ = 0.13%) in our independent dataset, consistent with the results obtained from an unweighted polygenic score (**Methods**).

Given the sex-dimorphism of fat distribution in humans, previously shown to have a genetic basis (5, 17), we next performed meta-analyses of WHRadjBMI in women and men separately (**Table 1** and **Supplementary Fig 5**). We found SNP-based heritability 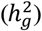 of WHRadjBMI, estimated using the restricted maximum likelihood method implemented in BOLT-REML (10) (**Methods**), to be stronger in women 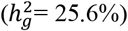 compared to men (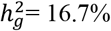, p_difference_ = 9 × 10^−85^; **Table 1, Supplementary Table 6**, and **Equation 2**). In addition to the heritability dimorphism, and in keeping with previous studies (5), we found signatures of sex-dimorphism amongst associated loci: a total of 266 loci associated with WHRadjBMI in women, compared to 91 loci in men (p < 5 × 10^−9^). Genome-wide, SNP effects on WHRadjBMI were strongly correlated between men and women (LD Score r_g_ = 0.514 (s.e. = 0.019), p = 3.43 × 10^−159^), but the consistency between the effect size of 266 female index SNPs on WHRadjBMI in women and men (adjusted R^2^ = 51%) was greater than the consistency between the effect size of 91 male index SNPs on WHRadjBMI in men and women (adjusted R^2^ = 9%). Of all associated index SNPs (p < 5 × 10^−9^ in the combined or sex-specific analyses), 105 SNPs were sex-dimorphic (p_diff_ < 3.3 × 10^−5^; (17) and **Methods**). Variants discovered in the combined sex analysis will be enriched for those with similar effects in each sex, while variants discovered in sex-specific analyses will be enriched for those with differing effects between sexes. In the absence of any sex-specific effects, we would only expect a slight shift towards stronger associations in women due to the larger available sample size in that analysis. However, we observed that of the 105 sex-dimorphic signals, 97 (92.4%) showed stronger effects in women compared to men (**Figure 1, Supplementary Fig 6**, and **Methods**). Scanning genome-wide for sex-dimorphic SNPs (p_diff_ < 5 × 10^−9^), regardless of their association p-values in the sex-specific analyses, we identified 61 sex-dimorphic SNPs after LD-based clumping (r^2^ < 0.05). Of these, 19 (31.1%) overlapped with the sex-dimorphic and genome-wide significant loci, and 54 (88.5%) had stronger effect in women than in men (**Supplementary Information**).

**Figure 1.**
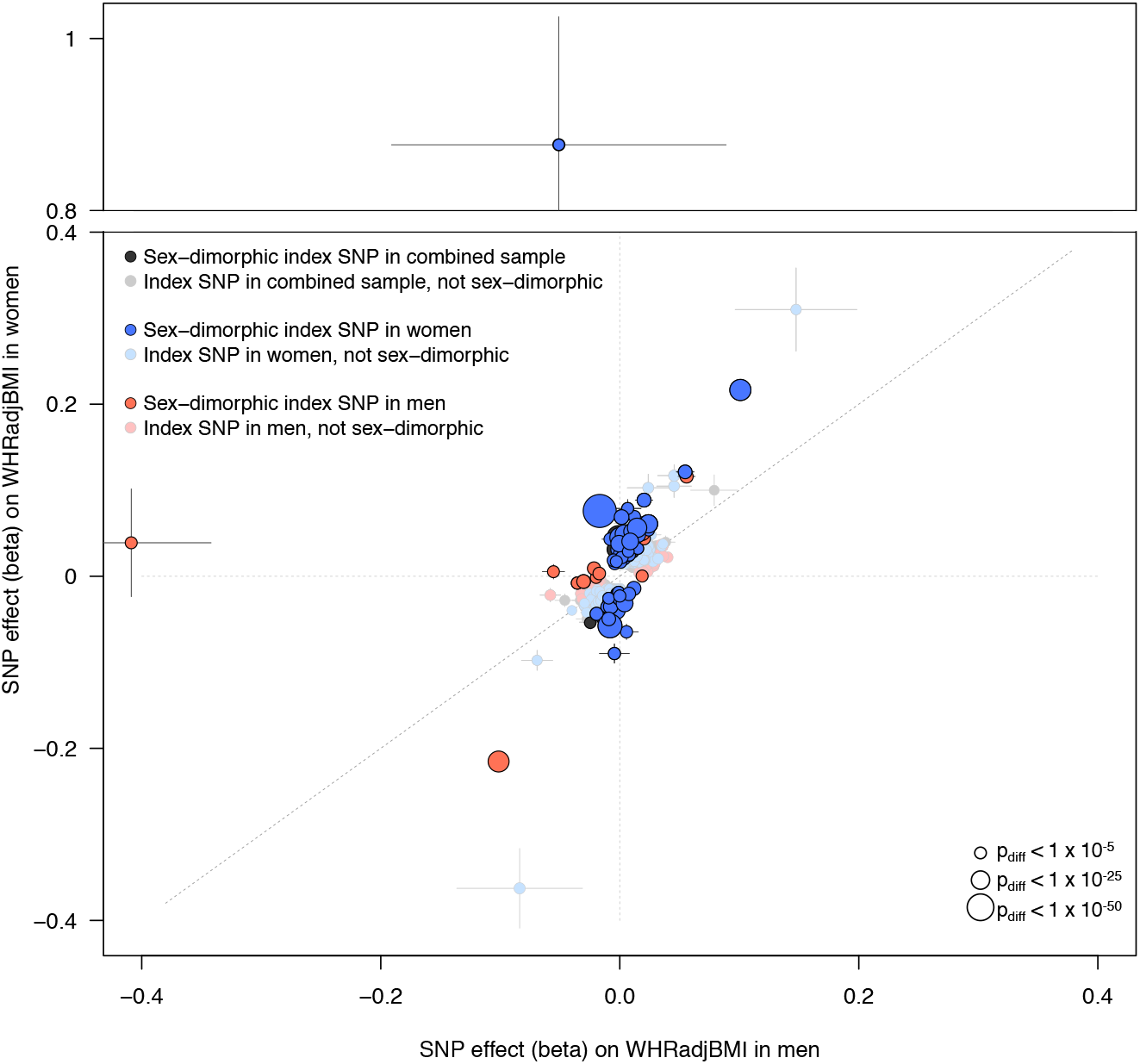
Sex-dimorphic association signals in fat distribution. For each associated locus from either the combined or sex-specific meta-analyses, we tested the index SNP for sex-dimorphism. We plot here all index SNPs from each of the three meta-analyses (combined, women only, and men only). SNPs that are significantly sex-dimorphic (p_diff_ < 3.3 × 10”^5^) are represented by boldly-colored circles, while index SNPs that are not sex-dimorphic are plotted with faded colors. Despite the expectation that SNPs identified in the combined sample (men and women, grey points) will be biased away from sex-dimorphism, and index SNPs identified in the sex-specific sample will be biased towards sex-dimorphism (due to winner’s curse), we observed stronger effects in women across all SNPs. Of the index SNPs from the men-only analysis (orange points), 14% showed evidence of sex-dimorphism. In contrast, ~29% of the index SNPs from the women-only analysis (blue points) show evidence of dimorphism. Over all sex-dimorphic SNPs, 92.4% show a stronger effect in women compared to men. Points are sized by the −log_10_(p_diff_) of the sex-dimorphism test. Horizontal bars indicate standard error in men; vertical bars indicate standard error in women.

Previous studies have shown that in addition to redistributing body fat, some WHRadjBMI variants are also associated with total body fat percentage (BF%) (5, 18–20). Of relevance to the biology of adipose tissue storage capacity, these studies have shown that these pleiotropic associations can occur in both directions: some alleles associated with higher WHRadjBMI are associated with higher total BF%, whilst others are associated with lower BF% (5, 18–20). To test the hypothesis that alleles associated with higher WHRadjBMI could have pleiotropic effects on total BF%, and that these effects could occur in both directions, we next investigated whether 346 index variants associated with WHRadjBMI also associated with BF%. Of the 59/346 variants associated with BF% in 443,001 European-ancestry UK Biobank individuals (p < 0.05/346 = 1.44 × 10^−4^), 25 SNPs associated with higher WHR and higher BF%, whilst 34 SNPs associated with higher WHR but lower BF% (**Figure 2**). These findings indicate that WHR-increasing alleles do not strictly influence BF% in one direction but rather can associate with either higher or lower BF%, yielding biological insight beyond the known epidemiological correlation between BF% and WHR. Additionally, a large proportion (29%) of WHRadjBMI index SNPs with a stronger effect in women had a BF% phenotype in men: 28 of the 97 female-specific WHRadjBMI SNPs were associated with BF% in men and 25 were associated with BF% in women (p < 0.05/105 = 4.8 × 10^−4^, **Supplementary Fig 7**). These variants appear to alter total BF% in men and women to a similar extent but distribute body fat between the upper and lower body to a much greater extent in women (**Supplementary Table 7-9** and **Supplementary Fig 7**). Finally, we tested the index SNPs from each of the meta-analyses (combined and sex-specific) in a recent GWAS of CT and MRI image-based measures of ectopic and subcutaneous fat depots (21). Adjusting for the three sample groups and the 8 depots examined in the imaging-based GWAS (p < 0.05/24 = 2.1 × 10^−3^), the alleles associated with higher WHRadjBMI were collectively associated with lower measures of subcutaneous fat, and higher measures of visceral fat, including pericardial and visceral adipose tissue (**Supplementary Fig 8**).

**Figure 2.**
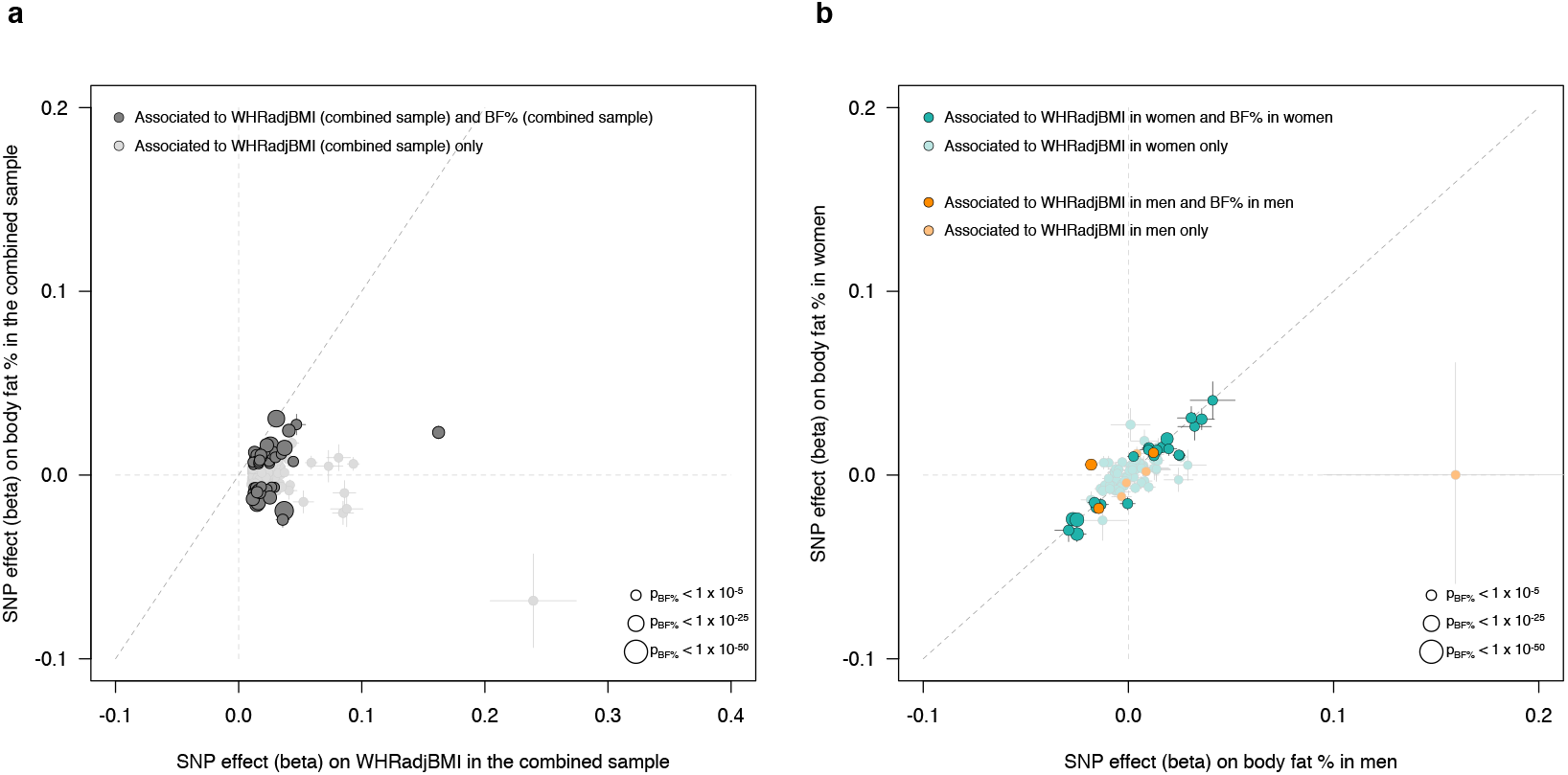
Effects of WHRadjBMI-associated SNPs on body fat percentage. (a) We investigated the impact of the 346 WHRadjBMI index SNPs (discovered in the combined analysis) on body fat percentage (BF%) in 449,001 UK Biobank individuals. Of the 346 SNPs, 59 (17.1%) are associated with BF% (p < 0.05/346 = 1.44 × 10^−4^, dark grey points). We oriented the effects of the SNPs to the WHRadjBMI-increasing effect, and found that 34 of the 59 BF%-associated SNPs associate with increased BF%, while 25/59 associate with decreased BF%, indicating that WHRadjBMI-associated SNPs can effect BF% in both directions, (b) Given the sex-dimorphic signature observed in WHRadjBMI-associated SNPs and the increased number of SNPs with stronger effects on WHRadjBMI in women we investigated the effect of the 105 sex-dimorphic index SNPs identified from the three meta-analyses (in the combined sample, in women only, in men only) on BF% in men or women separately. Of the 105 dimorphic SNPs, 97 were female specific (aquamarine points) and conferred a stronger effect on WHRadjBMI (on average) compared to the 8 male-specific SNPs (orange points). We plot the 105 sex-dimorphic SNPs by their effect on BF% in men (x-axis) and in women (y-axis). Of the 105 SNPs, 56 associate with BF% (p < 0.05/05 = 4.8 × 10^−3^). Despite the fact that these SNPs confer different effects on WHRadjBMI within sex-specific groups, we found that they confer relatively similar effects in BF% in sex-specific groups. All points are scaled in size to their strength of association in BF%.

## Discussion

In a meta-analysis of nearly 700,000 individuals, we have increased the number of loci associated to WHRadjBMI by more than seven-fold. Of all the detected signals, 105 are sex-dimorphic, consistent with previous findings (5). While we have performed the largest meta-analysis of a measure of body-fat distribution to date, a number of limitations remain. First, the substantially larger number of signals with a stronger effect in women compared to men may be influenced by the reduction in power (proportional to the product of sample size and SNP heritability) in the men-only analysis (**Table 1**) compared the women-only analysis. Despite the power difference in the sex-specific analyses, we would not expect the difference to result in 92% of signals conferring a stronger effect in women. Second, our replication sample was too small (~1% of the discovery) to formally replicate individual SNP associations, but the fact that 78% of the 300 previously unknown index associations showed consistent direction of effect suggests a low false positive rate. Finally, our meta-analysis focused only on European-ancestry samples. Given the very different body-fat distributions between people of European and non-European ancestry, and their very different risks of adiposity-related disease, studies in non-Europeans are urgently needed (22, 23).

In summary, the genetic variants and loci identified by this meta-analysis will likely provide starting points for further understanding the biology of body fat distribution and its relationship with disease.

## Materials and Methods

### I. Data and code availability

Code and data related to this project, including summary-level data from the meta-analyses, can be found online at https://github.com/lindgrengroup/fatdistnGW_AS.

### II. Phenotypes

To generate phenotypes for the waist-to-hip ratio (WHR) and waist-to-hip ratio adjusted for body mass index (WHRadjBMI) analyses in the UK Biobank data (**Supplementary Table 10**), we followed a phenotype conversion consistent with that performed in previous efforts investigating WHR and WHRadjBMI by the GIANT consortium (5, 24).

Using phenotype information from UK Biobank, we divided waist circumference by hip circumference to calculate the WHR measure, and then regressed the WHR measure on sex, age at assessment, age at assessment squared, and assessment centre. To generate the WHRadjBMI phenotype, we followed the same procedure and included body mass index (BMI) as an additional independent variable in the regression. We performed rank inverse normalization on the resulting residuals from the regression (**Supplementary Fig 9**) and used these normalized residuals as the tested phenotype in downstream genome-wide association testing. To generate phenotypes for the sex-specific analyses, we followed this same procedure but ran the regressions in sex-specific groups.

### III. Genome-wide association analyses

#### The UK Biobank data

We conducted genome-wide association testing in the second release (June 2017) version of the UK Biobank data(8); this release did not contain the corrected imputation at non-Haplotype Reference Consortium (HRC (25)) sites and we therefore subset all of the SNP data down to HRC SNPs only. The UK Biobank applied quality control to samples and genotypes and imputed the resulting genotype data using sequencing-based imputation reference panels. We performed all of our genome-wide association testing and downstream analyses on the publicly-available imputation data (released in bgen format).

We excluded samples as suggested by the UK Biobank upon release of the data (**Supplementary Table 11**). Sample exclusions included samples with genotype but no imputation information, samples with missingness > 5%, samples with mismatching phenotypic and genotypic sex, and samples that have withdrawn consent since the initiation of the project.

#### LD scores and genetic relationship matrix for BOLT-LMM

We implemented all genome-wide association studies (GWAS) in BOLT-LMM (10), which performs association testing using a linear mixed model. To run, BOLT-LMM requires three primary components: the (imputed) genotypic data for association testing; a reference panel of Linkage Disequilibrium (LD) scores per SNP, calculated using LD Score Regression (12); and genotype data used to approximate a genetic relationship matrix (GRM), which is the best method available in this sample size to account for all forms of relatedness, ancestral heterogeneity in the samples, and other (potentially hidden) structure in the data.

We performed sensitivity testing (**Supplementary Information, Supplementary Tables 12-13** and **Supplementary Fig 10**) using three LD Score reference datasets and four SNP-sets to construct the GRM. For our final GWAS, we used LD scores calculated from a randomly-selected, 9,748 unrelated UK Biobank samples (~2% of the full UK Biobank sample set; **Supplementary Information**) and a GRM constructed using: imputed SNPs with imputation info score > 0.8; MAF > 1%; Hardy Weinberg P-value > 1 × 10^−8^; genotype missingness < 1%, after converting imputed dosages to best-guess genotypes; LD pruned at a threshold (r^2^) of 0.2; and excluding the major histocompatibility complex, the lactase locus, and the inversions on chromosomes 8 and 17 (**Supplementary Information**).

#### Association testing

For genome-wide association testing, we used BOLT-LMM to run a linear mixed model (LMM). We tested SNPs with imputation quality (info) > 0.3, minor allele frequency (MAF) > 0.1% (equivalent to ~50 copies of the minor allele in the full sample), and only those single-nucleotide variants (SNVs) and single-nucleotide polymorphisms (SNPs) represented in the Haplotype Reference Consortium (25) imputation reference panel. We used only the standard LMM implementation (i.e., infinitesimal model, using --lmm) in BOLT-LMM (**Supplementary Fig 11–12**); we did not run association testing using a non-infinitesimal model. The only covariate used in the LMM was the SNP array used to genotype sample; we included no other covariates.

After association testing, we looked at known SNPs already reported in WHR, WHRadjBMI, and BMI (5, 24). At the previously-described loci, we checked correlation of frequency, beta, standard error, and −log10(p-value) between our UK Biobank GWAS and the previous GWAS results (**Supplementary Fig 13**). Additionally, we estimated genomic inflation (lambda) and the LD Score Intercept to check if the P-values were well calibrated (**Supplementary Table 2**); calculations were performed using the LD Score software (https://github.com/bulik/ldsc)

### IV. Meta-analysis of results from UK Biobank and GIANT

#### Data preparation and quality control

We downloaded summary-level results from previous meta-analyses of WHR and WHRadjBMI (https://portals.broadinstitute.org/collaboration/giant/index.php/GIANT_consortium_data_files and **Supplementary Information**) performed by the GIANT consortium (5). Marker names in both the GIANT data and UK Biobank were lifted over to their dbSNP151 identifier. We additionally renamed markers as “rsID:A1:A2” (where A1 was the tested allele in UK Biobank) to avoid ambiguity at multiallelic SNPs in the UK Biobank data. As the GIANT data was imputed with HapMap 2 (26, 27) data (hg18), we additionally lifted chromosomal positions to hg19 for this data. SNPs with a frequency difference > 15% between GIANT and UK Biobank were removed from the data (**Supplementary Fig 14**).

#### Meta-analysis and downstream quality control

We performed inverse variance-weighted fixed effects meta-analysis in METAL (28). To estimate LD score intercepts and genomic inflation (lambda) for the meta-analysis results, we first estimated LD scores from the same samples used to estimate the LD score reference for BOLT-LMM. LD scores were only estimated at high-quality SNPs (using the same criteria as used for SNPs included in the GRM in BOLT-LMM, but without applying a MAF threshold; **Supplementary Information**). We then calculated LD Score Regression intercepts and lambda with the LDSC software (12).

As an additional quality control check, we reran all of our GWAS using two different subsets of the UK Biobank samples: (1) the unrelated samples only, and (2) the unrelated white British samples only. These subsamples were selected to test if our initial UK Biobank-wide GWAS was confounded by either relatedness or ancestral heterogeneity. After running these GWAS, we meta-analyzed the results with the existing GIANT summary-level data and checked the concordance of our signals (**Supplementary Fig 2-3**).

### V. Identification of index and secondary signals

#### Linkage disequilibrium clumping

To identify genomic loci (i.e., genomic windows) containing independent association signals, we first constructed a reference dataset of best-guess genotypes from 20,275 unrelated UK Biobank samples (equivalent to 5% of the unrelated sample). We converted imputed dosages of SNPs with info score > 0.3 and MAF > 0.001% to best-guess genotypes using PLINK (version 1.9), (29, 30) and a conversion threshold (--hard-call-threshold) of 0.1 (**Supplementary Information**). SNPs with missingness > 5% after conversion or Hardy-Weinberg equilibrium p < 1 × 10^−7^ were removed.

We then used the PLINK ‘clumping’ algorithm to select top-associated SNPs (p < 5 × 10^−9^) and identify all SNPs in LD (r^2^ > 0.05) with the top associated SNP and ±5Mb away. We determined the genomic span of each LD-based clump and added 1kb up- and downstream as buffer to the region. If any of these windows overlapped, we merged them together into a single (larger) locus. As a sensitivity analysis, we ran clumping also using a smaller genomic window to calculate LD (±2Mb); the results were effectively unchanged, as <5 loci appeared independent using the ±2Mb window but were found to correlate using ±5Mb windows. Therefore, we report loci using the ±5Mb window.

#### Proximal conditional and joint testing

To identify index and secondary signals within each of the clumping-based loci, we ran proximal joint and conditional analysis as implemented in the Genome-wide Complex Trait Analysis (GCTA) software (31). We ran this model (--cojo-slct) using the summary-level data within each locus, the LD reference panel constructed from UK Biobank data and also used for the locus ‘clumping,’ and setting genome-wide significance with p < 5 × 10^−9^.

### VI. Validation in an independent dataset

We used an independent dataset EXTEND (7,721 individuals of European descent collected from South West England, **Supplementary Table 14**) to validate our findings. We extracted the index SNPs from the HRC imputed genotypes. To generate the WHRadjBMI variable, we regressed WHR on BMI, age, age-squared, sex and principal components 1-5. We then performed rank based inverse normalization on the resulting residuals. We validated the findings in 3 steps:

#### (1) Directional consistency

We checked for directional consistency between the effect of index SNPs on WHRadjBMI from the main meta-analysis and EXTEND. We performed linear regression of WHRadjBMI on each individual SNP. We ensured all alleles were aligned to the WHRadjBMI increasing allele in the original meta-analysis. We compared directions between all 346 index SNPs and then split these into novel and known signals to determine the number of novel signals showing consistent directionality.

#### (2) Variance explained

We evaluated the proportion of variance explained by including all the index SNPs into a linear regression model and calculated the adjusted *R*^2^. We performed these analyses using the lm() function in R.

#### (3) Polygenic scores

We created a weighted polygenic score based on the 346 index SNPs associated with WHRadjBMI. The weighted polygenic risk score (PRS) was calculated by summing the dosage of the WHRadjBMI-increasing alleles (weighted by the effect size on WHRadjBMI from the meta-analysis). We then performed linear regression to test the association between WHRadjBMI and the PRS in our independent dataset.

We sought to determine how likely the 5% of individuals carrying the most WHRadjBMI-increasing alleles were to meet the World Health Organization (WHO) WHR threshold used to diagnose metabolic syndrome (along with lipids and type 2 diabetes status) (13) compared to the 5% carrying the least. We used the WHR reference levels of > 0.9 in men and > 0.85 in women to define cases and WHR < 0.9 in men and < 0.85 in women to define controls (13). We excluded all individuals with missing data leaving a sample size of 7,513. We took 5% of individuals (7,513 × 0.05 = 376) from the two ends of weighted PRS and coded them as 1 or 2 respectively. We tested for the likelihood of the top 5% meeting the WHR threshold to diagnose metabolic syndrome (WHO criteria) compared to the bottom 5% using a binomial logistic regression model adjusting for age, age-squared, sex and principal components 1-5.

### VII. Collider bias analysis

Given that we had conditioned WHR on the BMI phenotype for analysis (and BMI and WHR are correlated; r = 0.433 in the UK Biobank data; **Supplementary Fig 15**), we tested all index signals found in the WHRadjBMI analysis for evidence of collider bias (15, 32). To do this, we ran meta-analyses of BMI and WHR using the UK Biobank samples and pre-existing summary-level data from GIANT (5, 24) (**Supplementary Methods**). We performed these meta-analyses using identical methods to the meta-analysis of WHRadjBMI.

Then, for each index SNP from the WHRadjBMI meta-analyses (combined as well as sex-specific) we extracted the association results from the BMI and WHR meta-analyses (**Supplementary Fig 4**). WHRadjBMI-associated SNPs with a stronger association for BMI than WHR show evidence of collider bias or pleiotropy. We additionally looked at the effect size and direction of effect in BMI and WHR, but whether the effects are from collider bias or pleiotropy cannot be determined from this data.

### VIII. Identification of sex-dimorphic signals

We estimated correlation between WHRadjBMI in females and in males using bivariate LD Score Regression analysis (12, 33).

We performed sex-specific GWAS in UK Biobank and meta-analyzed the results with publicly-available sex-specific data from the GIANT consortium. We identified the primary and secondary signals from these meta-analyses using methods identical to those performed in the combined analysis. We tested each primary and secondary signal for a sex-dimorphic effect by estimating the t-statistic:

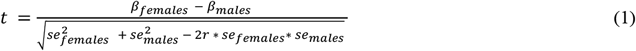

where *se* is the standard error and *r* is the genome-wide Spearman rank correlation coefficient between SNP effects in females and males. We estimated the t-statistic and the resulting so-called p_diff_ (p-value from a t-distribution with one degree of freedom (17)) as implemented in the EasyStrata software (34).

We tested a total of 2,162 different index SNPs for sex-dimorphism; we tested all of the secondary signals as well, but these signals are by definition in linkage disequilibrium with the index SNPs (and therefore not independent). Given that we tested for sex-dimorphism at index SNPs in not only WHRadjBMI but WHR and BMI as well, we performed a test at 1,502 distinct genomic loci. Therefore, we set significance for sex-dimorphism at a Bonferroni-corrected p = 0.05/1,502 = 3.3 × 10^−5^.

SNPs were determined to have a stronger effect in women if they fell into one of the following categories (abs, absolute value):

a. beta_females_ ≤ 0 and beta_males_ ≤ 0 and abs(beta_females_) > abs(beta_males_)
b. beta_females_ ≥ 0 and betamales ≥ 0 and abs(beta_females_) > abs(beta_males_)
c. beta_females_ ≤ 0 and beta_males_ ≥ 0 and pfemales < p_males_ and abs(beta_females_) > abs(beta_males_), or
d. beta_females_ ≥ 0 and beta_males_ ≤ 0 and p_females_ < p_males_ and abs(beta_females_) > abs(beta_males_)

### IX. Heritability calculations

#### SNP-based heritability calculations

We implemented all heritability calculations in BOLT-LMM.(10) We used the same genetic relationship matrix (GRM) to estimate SNP-based heritability as we did to run our GWAS (see *Genome-wide association analyses*). This GRM included 790,000 SNPs. Heritability was estimated using only the UK Biobank samples, for which we had individual level data; these estimates are likely more accurate than those resulting from only summary-level data. We used Restricted Maximum Likelihood Estimation, implemented as --reml in BOLT.

To test the impact of including lower-frequency SNPs in the heritability estimates, we constructed an additional GRM identically as we had for association testing but including no minor allele frequency threshold. This GRM included ~1.7M SNPs. Heritability analyses were calculated identically using this GRM and --reml in BOLT.

To calculate whether heritability estimates in men and women were sex-dimorphic, we used the following equation to generate a z-score:

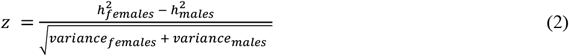

We then converted the z-scores to P-values using the following formula in the statistical programming language and software suite R (version 3.4):

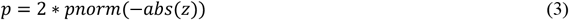

### X. Comparison of WHRadjBMI-associated SNPs in other fat distribution phenotypes

#### Comparison with body fat percentage

Similarly to Shungin et al (5), we carried out analysis on the 346 index SNPs and their association with BF% and WHR. We obtained association statistics for the 346 SNPs on BF% and WHR from a GWAS of 443,001 unrelated, European-ancestry UK Biobank individuals. We aligned all results to the WHR increasing allele and used a Bonferroni-corrected P-value (0.05/346 = 1..44 × 10^−4^) to determine if a SNP was associated with BF% (**Figure 2**). To determine whether sex-specific WHRadjBMI index SNPs have an adiposity phenotype, we took the 97 (female-specific) and 8 (male-specific) SNPs and independently compared their effects on WHRadjBMI and BF% in men and women. To identify which sex-dimorphic SNPs were strongly associated with BF% in men and women separately, we used a Bonferroni-corrected P-value of 0.05/105 (4.8 × 10^−4^) (**Supplementary Fig 7** and **Supplementary Table 9**). We obtained Pearson’s *r* correlations using the cor() function in R for each comparison.

#### Comparison with genome-wide analysis of depot-specific traits

Recently, Chu et al (21) performed a genome-wide association study of subcutaneous and ectopic fat depots, as measured by CT and MRI imaging, in a multi-ancestry sample. Since the meta-analysis results are publicly-available (https://grasp.nhlbi.nih.gov/FullResults.aspx and **Supplementary Information** for further details), we took the index SNPs from our WHRadjBMI meta-analyses (combined sample as well as sex-specific), checked for allele consistency, aligned effects to the reference allele, and tested for associations with the imaging based measures of subcutaneous and ectopic fat. We repeated these analyses in men and women separately. The depots investigated in the imaging-based GWAS were: pericardial tissue (PAT), PAT adjusted for height and weight (PATadjHtWt), subcutaneous adipose tissue (SAT), SAT Hounsfield units as measured by MRI (SATHU), visceral adipose tissue (VAT), VAT Hounsfield units (VATHU), ratio of VAT to SAT (VAT/SAT), and VAT adjusted for BMI (VATadjBMI).

We calculated Pearson’s r correlations between z-scores in WHRadjBMI (calculated by dividing the SNP beta by the standard error) and SNP z-scores reported in Chu et al (21). We evaluated significance of the correlation by performing a t-test (implemented as cor.test() in R). Correlations were considered significant if P-value < 0.05/3 sample groups/9 phenotypes = 1.9 × 10^−3^.

## Acknowledgements

This research was conducted using the UK Biobank Resource under Application Numbers 11867 and 9072. EXTEND data were provided by the Peninsula Research Bank, part of the NIHR Exeter Clinical Research Facility.

C.M.L is supported by the Li Ka Shing Foundation, WT-SSI/John Fell funds and by the NIHR Biomedical Research Centre, Oxford, by Widenlife and NIH (CRR00070 CR00.01).

S.L.P. is supported by a Veni Fellowship 016.186.071 (ZonMW) from the Dutch Organization for Scientific Research (Nederlandse Organisatie voor Wetenschappelijk Onderzoek, NWO).

H.Y. is funded by Diabetes UK RD Lawrence fellowship (grant: 17/0005594).

A.R.W. and T.M.F. are supported by the European Research Council grant: 323195:GLUCOSEGENES-FP7-IDEAS-ERC. R.B. is funded by the Wellcome Trust and Royal Society grant: 104150/Z/14/Z.

J.T. is funded by the ERDF and a Diabetes Research and Wellness Foundation Fellowship.

S.E.J. is funded by the Medical Research Council (grant: MR/M005070/1).

P.M.V. and J.Y. are funded by Australian National Health and Medical Research Council (1078037 and 1113400). J.Y. is supported by the Sylvia & Charles Viertel Charitable Foundation.

D.C.C.-C. is supported by a grant from the U.S. National Institutes of Health (K01 HL127265).

A.T.H. is a Wellcome Trust senior investigator and NIHR Senior Investigator.

## Authorship contributions

Data collection and analysis: S.L.P., C.S., A.P.M., A.R.W.,

Data interpretation: S.L.P., C.S., C.M.L., T.M.F., H.Y., A.P.M., C.G.

Study supervision: H.Y., T.F., S.L.P, C.M.L

First draft of the manuscript: S.L.P, C.M.L.

Critical revisions of the manuscript: all co-authors

## Abbreviations

BMI: Body mass index
WHR: Waist-to-hip ratio
WHRadjBMI: Waist-to-hip ratio, adjusted for body mass index
GWAS: Genome-wide association study
BF%: Body fat percentage
LD: Linkage disequilibrium
SNP: Single nucleotide polymorphism
UKBB: UK Biobank
T2D: Type 2 diabetes
GRS: Genetic risk score
CT: Computerized tomography
MR: Magnetic resonance imaging
BOLT-LMM: BOLT Linear Mixed Model
BOLT-REML: BOLT restricted maximum likelihood
PCA: Principal component analysis
SAT: Subcutaneous adipose tissue
VAT: Visceral adipose tissue
PAT: Pericardial adipose tissue
GRM: Genetic relationship matrix

